# Circulating electrolytes and the prevalence of atrial fibrillation and supraventricular ectopy: the Atherosclerosis Risk in Communities (ARIC) Study

**DOI:** 10.1101/645176

**Authors:** Alvaro Alonso, Mary R. Rooney, Lin Yee Chen, Faye L. Norby, Amy K. Saenger, Elsayed Z. Soliman, Wesley T. O’Neal, Katie C. Hootman, Elizabeth Selvin, Pamela L. Lutsey

**Affiliations:** Department of Epidemiology, Rollins School of Public Health, Emory University, Atlanta, GA; Division of Epidemiology and Community Health, School of Public Health, University of Minnesota, Minneapolis, MN; Cardiovascular Division, Department of Medicine, University of Minnesota Medical School, Minneapolis, MN; Chemistry Laboratory, Hennepin Healthcare, Minneapolis, MN; Epidemiological Cardiology Research Center (EPICARE), Department of Epidemiology and Prevention, Wake Forest School of Medicine, Winston-Salem, NC; Division of Cardiology, Department of Medicine, School of Medicine, Emory University, Atlanta, GA; Metabolic Research Unit, Clinical and Translational Science Center, Weill Cornell Medicine, New York, NY; Department of Epidemiology, Johns Hopkins Bloomberg School of Public Health, Baltimore, MD

## Abstract

**Background:** Evaluating associations of circulating electrolytes with atrial fibrillation (AF) and burden of supraventricular arrhythmias can give insights into arrhythmia pathogenesis.

**Methods:** We conducted a cross-sectional analysis of 6,398 participants of the Atherosclerosis Risk in Communities (ARIC) study, ages 71-90, with data on serum electrolytes (magnesium, calcium, potassium, phosphorus, chloride, sodium). Prevalence of AF was determined from study 12-lead electrocardiograms and prior history of AF-related hospitalizations. A subset of 317 participants also underwent electrocardiographic recordings for up to 14 days using the Zio^®^ patch. Burden of other supraventricular arrhythmias [premature atrial contractions (PACs), supraventricular tachycardia] was determined with the Zio^®^ patch. We used multivariable logistic and linear regression adjusting for potential confounders to determine associations of electrolytes with arrhythmia prevalence and burden.

**Results:** Among 6,394 eligible participants, 614 (10%) had prevalent AF. Participants in the top quintiles of magnesium [odds ratio (OR) 0.82, 95% confidence interval (CI) 0.62, 1.08], potassium (OR 0.82, 95%CI 0.68, 1.00), and phosphorus (OR 0.73, 95%CI 0.59, 0.89) had lower prevalence of AF compared to those in the bottom quintiles. No clear association was found for circulating chloride, calcium or sodium. Higher concentrations of circulating calcium were associated with lower prevalence of PACs using a standard 12-lead electrocardiogram, while higher concentrations of potassium, chloride and sodium were associated with higher PAC prevalence. Circulating electrolytes were not significantly associated with the burden of PACs or supraventricular tachycardia among a subset of 317 participants with extended electrocardiographic monitoring.

**Conclusion:** Concentrations of circulating electrolytes present complex associations with selected supraventricular arrhythmias. Future studies should evaluate underlying mechanisms.

## INTRODUCTION

Lower circulating magnesium and higher concentrations of phosphorus have been associated with increased risk of atrial fibrillation, a common arrhythmia (AF).^1–3^ Data on the association of other circulating electrolytes (such as calcium, chloride, sodium, and potassium) with the risk of AF are scarcer, though limited evidence suggests that low serum potassium and high chloride may be associated with increased AF risk.^4, 5^ Moreover, no prior research has explored the association of these circulating electrolytes with the burden of other supraventricular arrhythmias, including supraventricular tachycardia (SVT) or premature atrial contractions (PACs), in a community-based setting.

Given the role that electrolytes play in cardiac electrophysiology and prior evidence linking their circulating concentrations with AF risk, characterizing the association of electrolytes with supraventricular arrhythmias may refine our understanding of their contribution to atrial electrophysiology, which could inform, in turn, the development of preventive or treatment strategies for AF. To address this gap, we evaluated the cross-sectional associations of selected electrolytes with prevalence of AF and PACs in the Atherosclerosis Risk in Communities (ARIC) study. We also assessed the association of these electrolytes with the burden of supraventricular arrhythmias in a subset of ARIC study participants who underwent ambulatory continuous electrocardiographic monitoring.

## METHODS

### Study population

The ARIC study recruited 15,792 men and women 45 to 64 years of age from four communities in the United States (Forsyth County, North Carolina; Jackson, Mississippi; selected suburbs of Minneapolis, Minnesota; Washington County, Maryland) in 1987-89.^6^ Participants underwent follow-up visits in 1990-92, 1993-95, 1996-98, 2011-13, 2016-17 and 2018-19. During study visits, participants provided information on lifestyles and clinical variables, underwent a basic physical exam, including anthropometric and blood pressure measurements, and had blood samples collected after eight hours of fasting following standardized protocols. Participants also took part in follow-up phone calls, and were followed continuously for hospitalization and death. From 15,792 participants recruited by the ARIC cohort in 1987-89, 6,538 took part in ARIC visit 5 (68% of participants alive), and 6,394 had electrolyte data and met inclusion criteria (Figure 1). Of these, 317 individuals wore an electrocardiogram monitor for up to two weeks as part of an ancillary study. The Institutional Review Boards of all participating institutions approved the study protocol and participants provided written informed consent at baseline and during follow-up visits.

**Figure 1.**
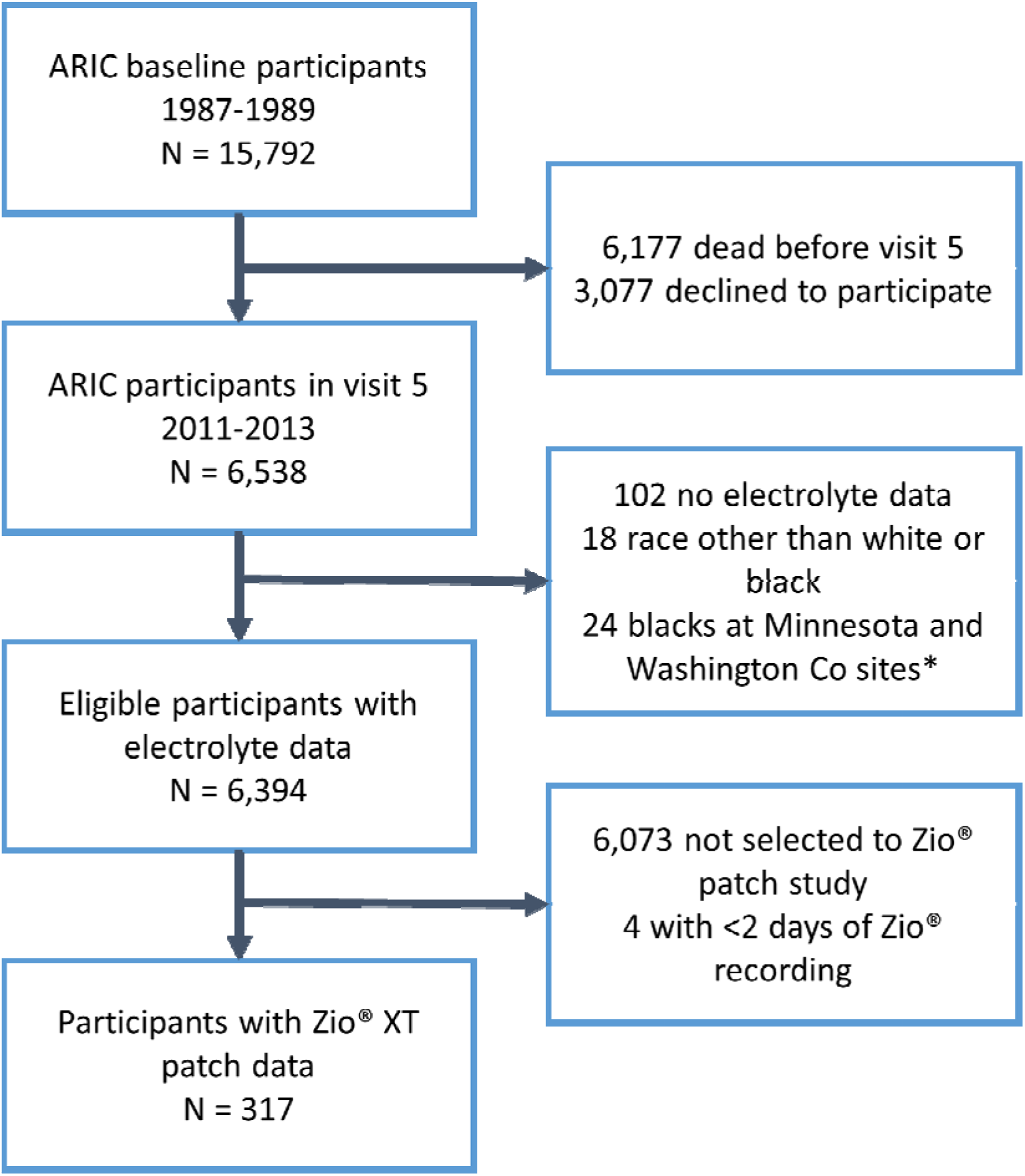
Flowchart of study participants, ARIC study. *Participants of race other than white or black, as well as blacks in the Minnesota and Washington County sites were excluded due to small numbers.

### Electrolyte measurements

Serum samples collected during the 2011-13 study exam (ARIC visit 5) had remained stored at −80°C in a central facility after collection and processing. Using these samples, in 2016 we measured magnesium, phosphorus, sodium, chloride, potassium, and calcium with a Roche COBAS 6000 chemistry analyzer (Roche Diagnostics, Indianapolis, IN). Magnesium was measured with a colorimetric (xylidyl blue) method, phosphorus with a photometric (molybdate) method, calcium with a colorimetric (NM-BAPTA) method, and sodium, chloride and potassium with an indirect ion selective electrode method. Coefficient of variation in measures from 242 duplicate samples were 1.9% for magnesium, 1.6% for phosphorus, 1.1% for calcium, 0.5% for sodium, 0.9% for chloride and 2.1% for potassium.

### Endpoint definitions

We defined prevalent AF at visit 5 based on (a) electrocardiographic evidence of the arrhythmia in any ARIC study exam or (b) presence of AF in any hospitalization discharge diagnosis occurring between baseline and ARIC visit 5 not associated with open cardiac surgery.^7^ At visit 5, a standard 10-second 12-lead electrocardiogram was used to define presence of PACs (Novacode code 1.0.S.1, sinus rhythm with ectopic supraventricular complexes).

A subset of 317 individuals participating in visit 5 wore the Zio^®^ Patch (iRhythm Technologies, Inc., San Francisco, CA), a leadless electrocardiogram monitor, for at least two days.^8^ The Zio^®^ Patch is a single-channel electrocardiographic monitor that provides up to two weeks of continuous electrocardiographic data. Data collected from the Zio^®^ Patch are processed using a proprietary algorithm [Zio ECG Utilization Service (ZEUS) System] that provides information on the number of PACs and SVT episodes during the wearing period.^9^ PAC burden was calculated as number of episodes per day by dividing the total number of PACs by the Zio^®^ Patch recording period (in days). We calculated SVT burden similarly.

### Measurement of other covariates

Participants self-reported information on age, sex, race, smoking, alcohol intake, and physical activity. Systolic and diastolic blood pressure, height and weight were measured using standard procedures. Prior use of medications was ascertained by asking participants to bring to the clinic visit any supplements or medications that they had used over the prior two weeks. Diabetes was defined as a fasting blood glucose ≥126 mg/dL, non-fasting glucose >200 mg/dL, use of antidiabetic medications, or self-reported history of a physician diagnosis of diabetes. History of coronary heart disease and heart failure was defined using self-reported information collected at the baseline visit and from adjudicated events during follow-up. Estimated glomerular filtration rate was calculated from measures of circulating creatinine and cystatin C.

### Statistical analysis

Before estimating associations between electrolytes and arrhythmias, we used multiple imputation with chained equations to avoid dropping observations due to missingness.^10^ This approach is more appropriate than dropping observations with missing values since it provides more precise estimates of the association and reduces the risk of selection bias.^11^ The imputation model included all the variables used as adjustment variables in multivariable models, listed below, as well as prevalent AF at visit 5 and the number of PACs on the 12-lead ECG. We created 20 imputed data sets using SAS PROC MI, performed separate analyses in each data set, and combined results using SAS PROC MIANALYZE. Of 6,394 eligible participants, 1,289 had missing data in at least one variable (mostly physical activity).

We assessed the association of circulating electrolytes with the prevalence of AF and presence of PACs in the visit 5 electrocardiogram using binary logistic regression, while we used multiple linear regression to assess associations with supraventricular arrhythmia burden as measured by the Zio^®^ patch. PAC and SVT burden were defined as total number of recorded episodes divided by the patch recording time (in days.) We also modeled the exposure variables as restricted cubic splines to more flexibly evaluate the continuous associations. Separate models were run for each electrolyte individually. We adjusted for age, sex, race, center, smoking, physical activity, alcohol intake, systolic and diastolic blood pressure, diuretic use, angiotensin converting enzyme inhibitor (ACEI) / angiotensin receptor blocker (ARB) use, other antihypertensive use, diabetes, heart failure, coronary heart disease, and estimated glomerular filtration rate. In a sensitivity analysis, we repeated the analysis excluding participants using diuretics, ACEIs or ARBs, since they have strong effects on the concentrations of many of the studied electrolytes.

## RESULTS

We included 6,394 eligible individuals with electrolyte data. Participants were 23% African-American, 59% female, and had a mean (standard deviation) age of 76 (5) years. Table 1 presents selected participant characteristics by sex and race.

**Table 1.** Selected participant characteristics by sex and race, ARIC study, 2011-2013

|  | White women | White men | African American women | African American men |
| --- | --- | --- | --- | --- |
| N | 2,761 | 2,171 | 987 | 475 |
| Age, years | 75.8 (5.3) | 76.3 (5.2) | 75.2 (5.3) | 74.8 (5.0) |
| Current smoker, % | 5.8 | 6.1 | 6.5 | 9.4 |
| Physical activity, score | 2.5 (0.8) | 2.8 (0.8) | 2.4 (0.7) | 2.6 (0.8) |
| Alcohol intake, g/week | 24.7 (48.2) | 50.5 (85.3) | 7.3 (26.1) | 27.6 (73.2) |
| Body mass index, kg/m<sup>2</sup> | 27.9 (5.7) | 28.5 (4.6) | 31.3 (7.2) | 28.9 (6.0) |
| Height, cm | 159 (6) | 174 (7) | 161 (6) | 175 (7) |
| Systolic blood pressure, mmHg | 130 (18) | 128 (17) | 137 (21) | 133 (19) |
| Diastolic blood pressure, mmHg | 66 (10) | 66 (11) | 70 (11) | 71 (11) |
| Antihypertensive medication |  |  |  |  |
| Diuretics, % | 30 | 25 | 41 | 36 |
| ACEI / ARB, % | 40 | 49 | 56 | 56 |
| Other antihypertensive, % | 14 | 16 | 14 | 13 |
| Diabetes, % | 27 | 34 | 44 | 45 |
| Heart failure, % | 8 | 11 | 14 | 16 |
| Coronary heart disease, % | 10 | 27 | 10 | 15 |
| eGFR, mL/min/1.73 m<sup>2</sup> | 64 (17) | 65 (18) | 67 (21) | 69 (21) |
| Atrial fibrillation, % | 8 | 14 | 5 | 5 |
| Premature atrial contraction, % | 12 | 12 | 19 | 16 |
| Magnesium, mg/dL | 2.0 (0.2) | 2.0 (0.2) | 2.0 (0.2) | 2.0 (0.2) |
| Calcium, mg/dL | 9.4 (0.4) | 9.3 (0.4) | 9.5 (0.5) | 9.3 (0.4) |
| Potassium, mmol/L | 4.0 (0.3) | 4.1 (0.3) | 4.0 (0.4) | 4.1 (0.4) |
| Phosphorus, mg/dL | 3.6 (0.4) | 3.3 (0.4) | 3.5 (0.5) | 3.2 (0.5) |
| Chloride, mmol/L | 100 (3) | 101 (3) | 101 (3) | 101 (3) |
| Sodium, mmol/L | 139 (3) | 139 (2) | 140 (3) | 139 (2) |

Values correspond to mean (SD) or %. Based on imputed dataset. ACEI: angiotensin converting enzyme
inhibitor. ARB: angiotensin receptor blocker. eGFR: estimated glomerular filtration rate.

Prevalent AF was present in 614 participants (10% prevalence; 199 with evidence of AF in at least a study ECG and 415 exclusively detected from prior hospitalizations). The associations of prevalent AF with circulating electrolytes modeled using restricted cubic splines are shown in Figure 2. Overall, associations were non-linear for most electrolytes and, therefore, we categorized them in approximate quintiles. Table 2 reports odds ratios (OR) and 95% confidence intervals (95%CI) of prevalent AF by quintiles of circulating electrolytes. In multivariable analyses, participants in the top quintiles of magnesium (OR 0.82, 95%CI 0.62, 1.08), potassium (OR 0.82, 95%CI 0.68, 1.00), and phosphorus (OR 0.73, 95%CI 0.59, 0.89) had lower prevalence of AF compared to those in the bottom quintiles. No clear association was found for circulating chloride, calcium or sodium. The inverse association of circulating magnesium and phosphorus with prevalence of AF remained in 2516 participants not using diuretics, ACEIs, or ARBs (Supplemental Table S1).

**Table 2.**
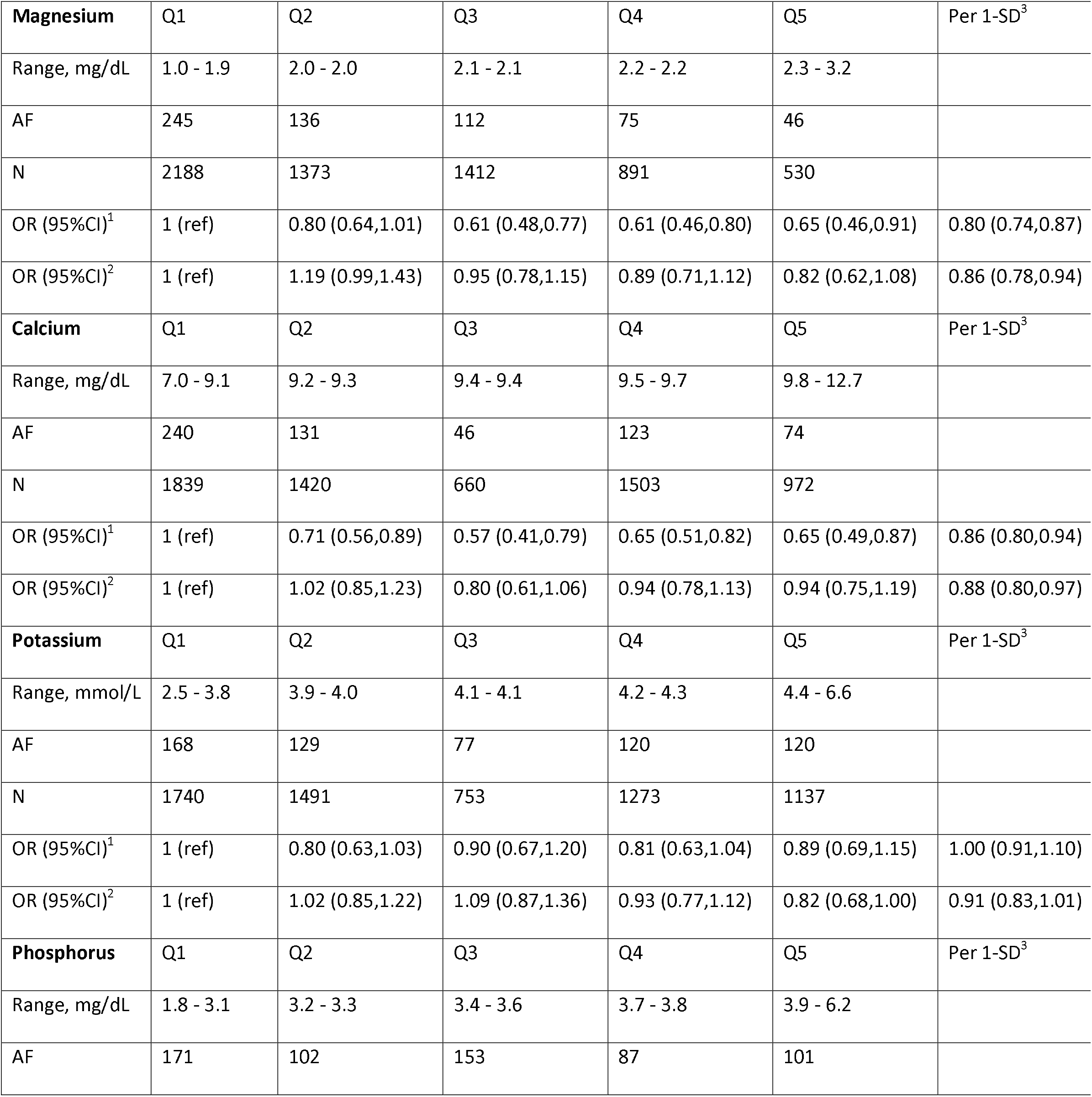
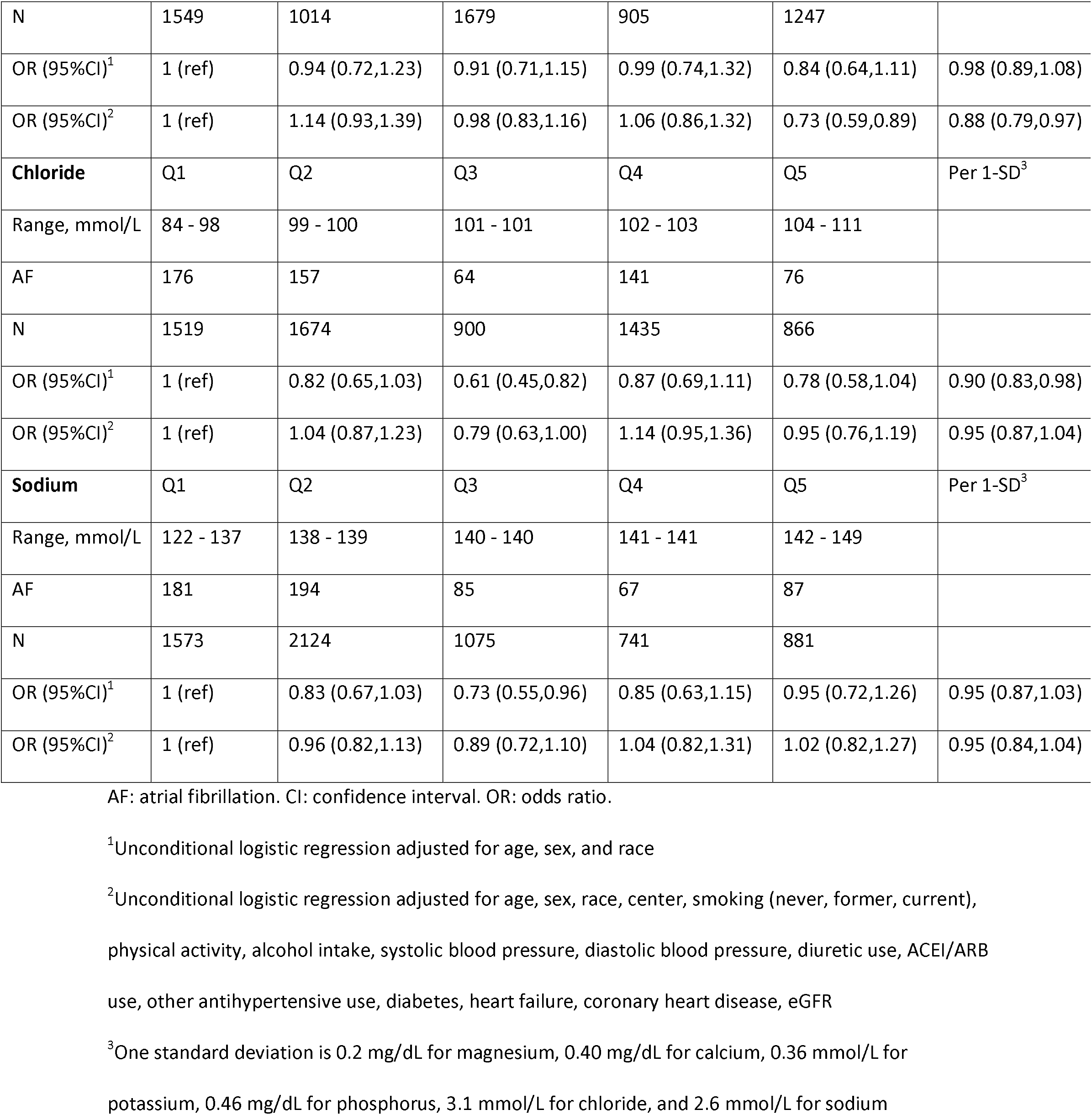
Association of circulating electrolytes with prevalence of atrial fibrillation, ARIC study, 2011-2013

|  |  |  |  |  |  |  |
| --- | --- | --- | --- | --- | --- | --- |
| **Magnesium** | Q1 | Q2 | Q3 | Q4 | Q5 | Per 1-SD<sup>3</sup> |
| Range, mg/dL | 1.0 - 1.9 | 2.0 - 2.0 | 2.1 - 2.1 | 2.2 - 2.2 | 2.3 - 3.2 |  |
| AF | 245 | 136 | 112 | 75 | 46 |  |
| N | 2188 | 1373 | 1412 | 891 | 530 |  |
| OR (95%CI)<sup>1</sup> | 1 (ref) | 0.80 (0.64,1.01) | 0.61 (0.48,0.77) | 0.61 (0.46,0.80) | 0.65 (0.46,0.91) | 0.80 (0.74,0.87) |
| OR (95%CI)<sup>2</sup> | 1 (ref) | 1.19 (0.99,1.43) | 0.95 (0.78,1.15) | 0.89 (0.71,1.12) | 0.82 (0.62,1.08) | 0.86 (0.78,0.94) |
| **Calcium** | Q1 | Q2 | Q3 | Q4 | Q5 | Per 1-SD<sup>3</sup> |
| Range, mg/dL | 7.0 - 9.1 | 9.2 - 9.3 | 9.4 - 9.4 | 9.5 - 9.7 | 9.8 - 12.7 |  |
| AF | 240 | 131 | 46 | 123 | 74 |  |
| N | 1839 | 1420 | 660 | 1503 | 972 |  |
| OR (95%CI)<sup>1</sup> | 1 (ref) | 0.71 (0.56,0.89) | 0.57 (0.41,0.79) | 0.65 (0.51,0.82) | 0.65 (0.49,0.87) | 0.86 (0.80,0.94) |
| OR (95%CI)<sup>2</sup> | 1 (ref) | 1.02 (0.85,1.23) | 0.80 (0.61,1.06) | 0.94 (0.78,1.13) | 0.94 (0.75,1.19) | 0.88 (0.80,0.97) |
| **Potassium** | Q1 | Q2 | Q3 | Q4 | Q5 | Per 1-SD<sup>3</sup> |
| Range, mmol/L | 2.5 - 3.8 | 3.9 - 4.0 | 4.1 - 4.1 | 4.2 - 4.3 | 4.4 - 6.6 |  |
| AF | 168 | 129 | 77 | 120 | 120 |  |
| N | 1740 | 1491 | 753 | 1273 | 1137 |  |
| OR (95%CI)<sup>1</sup> | 1 (ref) | 0.80 (0.63,1.03) | 0.90 (0.67,1.20) | 0.81 (0.63,1.04) | 0.89 (0.69,1.15) | 1.00 (0.91,1.10) |
| OR (95%CI)<sup>2</sup> | 1 (ref) | 1.02 (0.85,1.22) | 1.09 (0.87,1.36) | 0.93 (0.77,1.12) | 0.82 (0.68,1.00) | 0.91 (0.83,1.01) |
| **Phosphorus** | Q1 | Q2 | Q3 | Q4 | Q5 | Per 1-SD<sup>3</sup> |
| Range, mg/dL | 1.8 - 3.1 | 3.2 - 3.3 | 3.4 - 3.6 | 3.7 - 3.8 | 3.9 - 6.2 |  |
| AF | 171 | 102 | 153 | 87 | 101 |  |
| N | 1549 | 1014 | 1679 | 905 | 1247 |  |
| OR (95%CI)<sup>1</sup> | 1 (ref) | 0.94 (0.72,1.23) | 0.91 (0.71,1.15) | 0.99 (0.74,1.32) | 0.84 (0.64,1.11) | 0.98 (0.89,1.08) |
| OR (95%CI)<sup>2</sup> | 1 (ref) | 1.14 (0.93,1.39) | 0.98 (0.83,1.16) | 1.06 (0.86,1.32) | 0.73 (0.59,0.89) | 0.88 (0.79,0.97) |
| **Chloride** | Q1 | Q2 | Q3 | Q4 | Q5 | Per 1-SD<sup>3</sup> |
| Range, mmol/L | 84 - 98 | 99 - 100 | 101 - 101 | 102 - 103 | 104 - 111 |  |
| AF | 176 | 157 | 64 | 141 | 76 |  |
| N | 1519 | 1674 | 900 | 1435 | 866 |  |
| OR (95%CI)<sup>1</sup> | 1 (ref) | 0.82 (0.65,1.03) | 0.61 (0.45,0.82) | 0.87 (0.69,1.11) | 0.78 (0.58,1.04) | 0.90 (0.83,0.98) |
| OR (95%CI)<sup>2</sup> | 1 (ref) | 1.04 (0.87,1.23) | 0.79 (0.63,1.00) | 1.14 (0.95,1.36) | 0.95 (0.76,1.19) | 0.95 (0.87,1.04) |
| **Sodium** | Q1 | Q2 | Q3 | Q4 | Q5 | Per 1-SD<sup>3</sup> |
| Range, mmol/L | 122 - 137 | 138 - 139 | 140 - 140 | 141 - 141 | 142 - 149 |  |
| AF | 181 | 194 | 85 | 67 | 87 |  |
| N | 1573 | 2124 | 1075 | 741 | 881 |  |
| OR (95%CI)<sup>1</sup> | 1 (ref) | 0.83 (0.67,1.03) | 0.73 (0.55,0.96) | 0.85 (0.63,1.15) | 0.95 (0.72,1.26) | 0.95 (0.87,1.03) |
| OR (95%CI)<sup>2</sup> | 1 (ref) | 0.96 (0.82,1.13) | 0.89 (0.72,1.10) | 1.04 (0.82,1.31) | 1.02 (0.82,1.27) | 0.95 (0.84,1.04) |

AF: atrial fibrillation. CI: confidence interval. OR: odds ratio.
<sup>1</sup>Unconditional logistic regression adjusted for age, sex, and race
<sup>2</sup>Unconditional logistic regression adjusted for age, sex, race, center, smoking (never, former, current), physical activity, alcohol intake, systolic blood pressure, diastolic blood pressure, diuretic use, ACEI/ARB use, other antihypertensive use, diabetes, heart failure, coronary heart disease, eGFR
<sup>3</sup>One standard deviation is 0.2 mg/dL for magnesium, 0.40 mg/dL for calcium, 0.36 mmol/L for potassium, 0.46 mg/dL for phosphorus, 3.1 mmol/L for chloride, and 2.6 mmol/L for sodium

**Figure 2.**
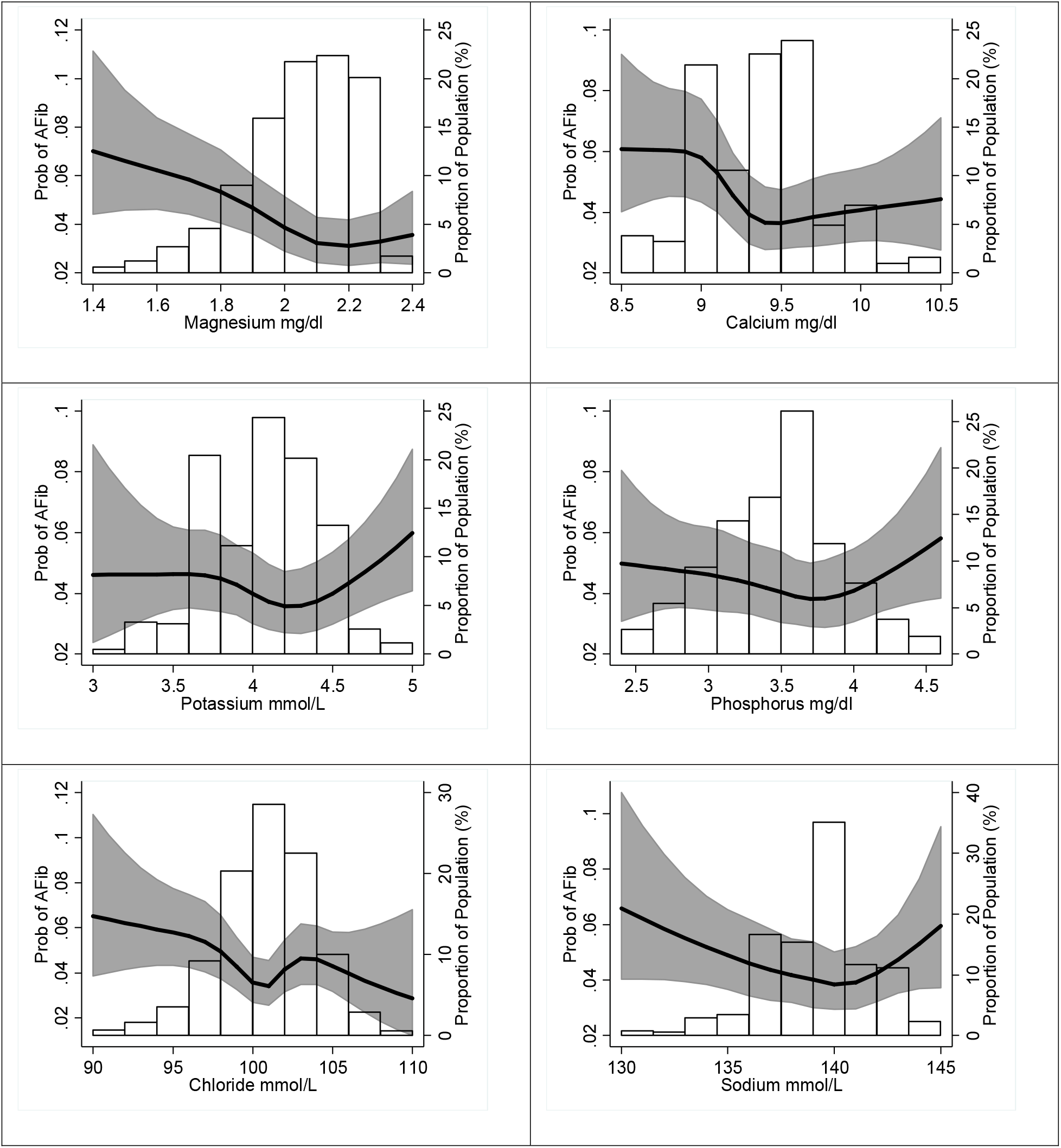
Association of the concentration of circulating electrolytes with the prevalence of AF, ARIC study, 2011-2013. Predicted probabilities calculated from logistic regression modeling circulating electrolytes with restricted cubic splines and adjusting for age, sex, and race. Values below the 1^st^ percentile and above the 99^th^ percentile were removed for interpretability.

Among 5,722 eligible participants without prevalent AF, wandering atrial pacemaker, complete atrioventricular block, Wolf-Parkinson-White pattern, or pacemaker, 748 (13%) had at least one PAC in the standard 12-lead electrocardiogram. Table 3 shows the associations of circulating electrolytes with PAC prevalence. Higher concentrations of circulating calcium were associated with lower prevalence of PACs (OR 0.76, 95%CI 0.63, 0.92 comparing highest to lowest categories). In contrast, higher concentrations of chloride and sodium were associated with higher PAC prevalence (OR 1.16, 95%CI 0.96, 1.39, and OR 1.20, 95%CI 1.00, 1.43, respectively, comparing highest to lowest categories). No associations were observed with circulating magnesium, potassium or phosphorus. In the 2,516 participants not using diuretics, ACEIs or ARBs, associations with PAC prevalence were strongly attenuated (Supplemental Table S2).

**Table 3.** Association of circulating electrolytes with prevalence of premature atrial contractions in standard 12-lead electrocardiogram, ARIC study, 2011-2013

| **Magnesium** | Q1 | Q2 | Q3 | Q4 | Q5 | Per 1-SD<sup>3</sup> |
| --- | --- | --- | --- | --- | --- | --- |
| Range, mg/dL | 1.0 - 1.9 | 2.0 - 2.0 | 2.1 - 2.1 | 2.2 - 2.2 | 2.3 - 3.2 |  |
| PAC present | 256 | 152 | 158 | 119 | 63 |  |
| N | 1920 | 1233 | 1284 | 811 | 474 |  |
| OR (95%CI)<sup>1</sup> | 1 (ref) | 0.94 (0.75,1.17) | 0.97 (0.78,1.21) | 1.13 (0.89,1.44) | 0.97 (0.71,1.31) | 1.00 (0.92,1.08) |
| OR (95%CI)<sup>2</sup> | 1 (ref) | 0.95 (0.80,1.11) | 1.03 (0.88,1.22) | 1.15 (0.96,1.38) | 0.97 (0.77,1.23) | 1.05 (0.96,1.14) |
| **Calcium** | Q1 | Q2 | Q3 | Q4 | Q5 | Per 1-SD<sup>3</sup> |
| Range, mg/dL | 7.0 - 9.1 | 9.2 - 9.3 | 9.4 - 9.4 | 9.5 - 9.7 | 9.8 - 12.7 |  |
| PAC present | 233 | 180 | 75 | 161 | 99 |  |
| N | 1580 | 1276 | 608 | 1368 | 890 |  |
| OR (95%CI)<sup>1</sup> | 1 (ref) | 0.98 (0.79,1.22) | 0.83 (0.62,1.10) | 0.75 (0.60,0.93) | 0.67 (0.52,0.87) | 0.89 (0.82,0.96) |
| OR (95%CI)<sup>2</sup> | 1 (ref) | 1.20 (1.02,1.40) | 1.02 (0.82,1.26) | 0.89 (0.76,1.04) | 0.76 (0.63,0.92) | 0.87 (0.80, 0.94) |
| **Potassium** | Q1 | Q2 | Q3 | Q4 | Q5 | Per 1-SD<sup>3</sup> |
| Range, mmol/L | 2.5 - 3.8 | 3.9 - 4.0 | 4.1 - 4.1 | 4.2 - 4.3 | 4.4 - 6.5 |  |
| PAC present | 188 | 150 | 95 | 157 | 158 |  |
| N | 1558 | 1351 | 669 | 1144 | 1000 |  |
| OR (95%CI)<sup>1</sup> | 1 (ref) | 0.94 (0.74,1.19) | 1.26 (0.96,1.65) | 1.24 (0.98,1.57) | 1.36 (1.08,1.73) | 1.17 (1.07, 1.27) |
| OR (95%CI)<sup>2</sup> | 1 (ref) | 0.86 (0.73,1.01) | 1.15 (0.95,1.40) | 1.09 (0.93,1.28) | 1.07 (0.90,1.27) | 1.12 (1.02,1.23) |
| **Phosphorus** | Q1 | Q2 | Q3 | Q4 | Q5 | Per 1-SD<sup>3</sup> |
| Range, mg/dL | 1.8 - 3.1 | 3.2 - 3.3 | 3.4 - 3.6 | 3.7 - 3.8 | 3.9 - 6.2 |  |
| PAC present | 199 | 118 | 190 | 94 | 147 |  |
| N | 1360 | 908 | 1514 | 810 | 1130 |  |
| OR (95%CI)<sup>1</sup> | 1 (ref) | 0.88 (0.68,1.13) | 0.87 (0.69,1.08) | 0.81 (0.62,1.07) | 0.94 (0.73,1.20) | 0.93 (0.85,1.02) |
| OR (95%CI)<sup>2</sup> | 1 (ref) | 1.00 (0.84,1.20) | 0.97 (0.84,1.13) | 0.90 (0.74,1.09) | 0.98 (0.82,1.15) | 0.89 (0.81,0.98) |
| **Chloride** | Q1 | Q2 | Q3 | Q4 | Q5 | Per 1-SD<sup>3</sup> |
| Range, mmol/L | 84 - 98 | 99 - 100 | 101 - 101 | 102 - 103 | 104 - 110 |  |
| PAC present | 176 | 189 | 86 | 179 | 118 |  |
| N | 1328 | 1502 | 827 | 1284 | 781 |  |
| OR (95%CI)<sup>1</sup> | 1 (ref) | 0.95 (0.76,1.19) | 0.76 (0.57,1.00) | 1.08 (0.86,1.36) | 1.11 (0.85,1.44) | 1.05 (0.97,1.13) |
| OR (95%CI)<sup>2</sup> | 1 (ref) | 0.98 (0.84,1.14) | 0.79 (0.65,0.96) | 1.15 (0.99,1.34) | 1.16 (0.96,1.39) | 1.08 (0.99,1.17) |
| **Sodium** | Q1 | Q2 | Q3 | Q4 | Q5 | Per 1-SD<sup>3</sup> |
| Range, mmol/L | 122 - 137 | 138 - 139 | 140 - 140 | 141 - 141 | 142 - 149 |  |
| PAC present | 162 | 233 | 130 | 89 | 134 |  |
| N | 1376 | 1913 | 985 | 664 | 784 |  |
| OR (95%CI)<sup>1</sup> | 1 (ref) | 1.07 (0.86,1.33) | 1.14 (0.89,1.47) | 1.13 (0.85,1.50) | 1.41 (1.09,1.82) | 1.11 (1.03,1.20) |
| OR (95%CI)<sup>2</sup> | 1 (ref) | 0.95 (0.83,1.09) | 1.02 (0.86,1.21) | 1.00 (0.82,1.22) | 1.20 (1.00,1.43) | 1.11 (1.02,1.20) |

CI: confidence interval. OR: odds ratio. PAC: premature atrial contraction
<sup>1</sup>Unconditional logistic regression adjusted for age, sex, and race
<sup>2</sup>Unconditional logistic regression adjusted for age, sex, race, center, smoking (never, former, current), physical activity, alcohol intake, systolic blood pressure, diastolic blood pressure, diuretic use, ACEI/ARB use, other antihypertensive use, diabetes, heart failure, coronary heart disease, eGFR
<sup>3</sup>One standard deviation is 0.2 mg/dL for magnesium, 0.40 mg/dL for calcium, 0.36 mmol/L for potassium, 0.46 mg/dL for phosphorus, 3.1 mmol/L for chloride, and 2.6 mmol/L for sodium

Among the subset of 317 ARIC participants who underwent extended electrocardiographic recording with the Zio^®^ patch for at least two days, 278 did not have AF. Supplemental Table S3 includes descriptive information of the participants who wore the patch. They were all white, slightly older and with a similar prevalence of comorbidities than participants not selected to wear the patch. Mean wearing time was 13.1 days and mean analyzable time was 12.7 days. All 278 participants without AF presented at least one PAC during their recording, and the mean (median) PAC burden in counts per day was 1,207 (236). None of the circulating electrolytes, categorized either in approximate tertiles or as continuous variables, were associated with PAC burden (Supplemental Table S4).

Prevalence of at least a single episode of SVT was 91% (253 participants out of 278), with a mean of 3 episodes per day among those with at least one recorded episode. Circulating electrolytes were not associated with presence of SVT in the Zio^®^ patch (Supplemental Table S5) or with SVT burden (Supplemental Table S6). Figure 3 summarizes the associations across electrolytes and endpoints.

**Figure 3.**
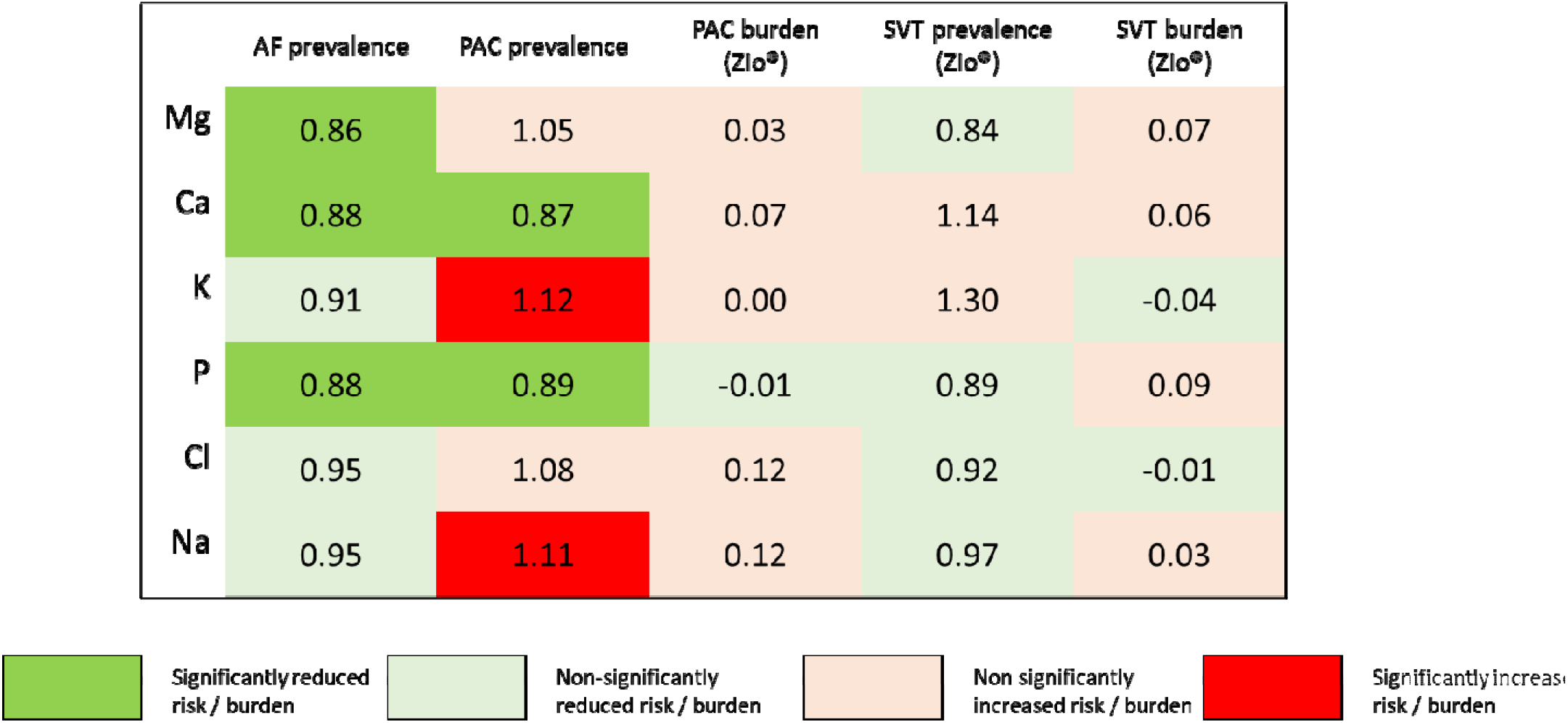
Summary of associations of circulating electrolytes with prevalence and burden of atrial fibrillation (AF), premature atrial contractions (PAC), supraventricular tachycardia (SVT). Columns for prevalence report odds ratios per one standard deviation increase; columns for burden present differences in episodes per day per one standard deviation increase. Green colors correspond to lower prevalence or burden, while red or beige colors correspond to higher prevalence or burden. One standard deviation is 0.2 mg/dL for magnesium, 0.40 mg/dL for calcium, 0.36 mmol/L for potassium, 0.46 mg/dL for phosphorus, 3.1 mmol/L for chloride, and 2.6 mmol/L for sodium.

## DISCUSSION

In this community-based sample of older adults, we observed a lower prevalence of AF among persons with higher concentrations of circulating magnesium, potassium, phosphorus, and chloride. Likewise, higher concentrations of calcium were associated with lower prevalence of PACs in a standard 12-lead electrocardiogram, while higher of chloride and sodium were associated with higher PAC prevalence. In contrast, circulating electrolytes were not associated with PAC or SVT burden in a small subset of participants with extended heart rhythm monitoring, but we did not have a sufficient sample size to robustly explore this research question.

These findings complement the existing literature on magnesium and supraventricular arrhythmias. Several prospective studies, including our prior work in the ARIC cohort, have reported lower risk of AF among individuals with higher concentrations of circulating magnesium.^1, 2^ Oral magnesium supplementation reduces the risk of postoperative AF,^12^ and a Mendelian randomization analysis reported an association of higher genetically-determined circulating magnesium with lower AF risk.^13^ These studies offer robust support for a potential causal effect of magnesium on the incidence of this arrhythmia. However, we did not observe associations between circulating magnesium and PAC presence or burden. PACs are associated with increased AF risk and are considered a potential endophenotype of AF.^14^ The lack of association in our analysis is consistent with a small randomized study reporting no effect of oral magnesium supplementation on PAC burden.^15^ As a whole, these findings suggest that magnesium may reduce AF risk through pathways other than facilitation of atrial ectopy. Future studies should test the efficacy of magnesium supplementation for prevention of AF and explore the underlying pathophysiological mechanisms.

The evidence on associations between other electrolytes and supraventricular arrhythmias in large community-based studies is sparse. In the Rotterdam study, hypokalemia (serum potassium <3.5 mmol/L) was associated with increased risk of AF compared to normokalemia.^4^ This finding is consistent with our results, in which prevalence of AF was lower among individuals with higher circulating potassium. Severe hypokalemia is associated with cardiac arrhythmias and affects atrial electrical activity in diverse ways.^16^ Higher concentrations of circulating phosphorus were associated with lower prevalence of AF in the present study. These results are unexpected and contrary to a previous prospective analysis of the ARIC cohort, which reported higher risk of AF associated with elevated circulating phosphorus.^3^ Differences in study design and the age of the population being studied could explain the inconsistent findings. Similar to this cross-sectional analysis, circulating calcium was not associated with incident AF in a previous analysis of the ARIC cohort.^3^ Calcium, however, plays a key role in cardiac electrophysiology and alterations in intracellular handling of calcium is an established arrhythmogenic mechanism.^17^ Whether changes in circulating calcium have any effect on these mechanisms remains unknown. No prior studies have evaluated the association of circulating chloride or sodium with AF risk or PAC prevalence. At the same time, electrophysiologic effects of variation in concentrations of sodium and chloride within the normal range are not well described. Thus, the significance of our findings regarding these two electrolytes—lower prevalence of AF associated with higher chloride and higher prevalence of PACs with higher sodium—is unclear. Future studies should aim to replicate these observations.

This study has notable strengths, including the relatively large sample size, the inclusion of a community-based sample, and detailed information on potential major confounders. There were also a number of limitations. Our study has potentially limited generalizability to younger individuals. The cross-sectional study design precludes establishing temporality of the associations and we cannot rule out the possibility of reverse causation. Finally, the number of participants undergoing long-term electrocardiographic monitoring was small, limiting the evaluation of associations in this group.

In summary, we observed that concentrations of circulating electrolytes present complex associations with selected supraventricular arrhythmias. These findings provide the rationale for future studies exploring the mechanisms underlying these associations and testing interventions for the prevention of arrhythmias by modifying circulating electrolytes.

## Supporting information

Supplementary Results

## ACKNOWLEDGMENTS

The authors thank the staff and participants of the ARIC study for their important contributions.

## REFERENCES

1. Misialek JR, Lopez FL, Lutsey PL, et al. Serum and dietary magnesium and incidence of atrial fibrillation in whites and in African Americans--Atherosclerosis Risk in Communities (ARIC) Study. Circ J. 2013;77:323–329.

2. Khan AM, Lubitz SA, Sullivan LM, et al. Low serum magnesium and the development of atrial fibrillation in the community: the Framingham Heart Study. Circulation. 2013;127:33–38.

3. Lopez FL, Agarwal SK, Grams ME, et al. Relation of serum phosphorus levels to the incidence of atrial fibrillation (from the Atherosclerosis Risk In Communities [ARIC] Study). Am J Cardiol. 2013;111:857–862.

4. Krijthe BP, Heeringa J, Kors JA, et al. Serum potassium levels and the risk of atrial fibrillation: the Rotterdam Study. Int J Cardiol. 2013;168:5411–5415.

5. Svagzdiene M, Sirvinskas E. Changes in serum electrolyte levels and their influence on the incidence of atrial fibrillation after coronary artery bypass grafting surgery. Medicina (Kaunas). 2006;42:208–214.

6. The ARIC Investigators. The Atherosclerosis Risk in Communities (ARIC) study: design and objectives. Am J Epidemiol. 1989;129:687–702.

7. Alonso A, Agarwal SK, Soliman EZ, et al. Incidence of atrial fibrillation in whites and African-Americans: the Atherosclerosis Risk in Communities (ARIC) study. Am Heart J. 2009;158:111–117.

8. Chen LY, Agarwal SK, Norby FL, et al. Persistent but not paroxysmal atrial fibrillation is independently associated with lower cognitive function: the Atherosclerosis Risk in Communities (ARIC) Study. J Am Coll Cardiol. 2016;67:1379–1380.

9. Turakhia MP, Ullal AJ, Hoang DD, et al. Feasibility of extended ambulatory electrocardiogram monitoring to identify silent atrial fibrillation in high-risk patients: the Screening Study for Undiagnosed Atrial Fibrillation (STUDY-AF). Clin Cardiol. 2015;38:285–292.

10. White IR, Royston P, Wood AM. Multiple imputation using chained equations: issues and guidance for practice. Stat Med. 2011;30:377–399.

11. Klebanoff MA, Cole SR. Use of multiple imputation in the epidemiologic literature. Am J Epidemiol. 2008;168:355–357.

12. Arsenault KA, Yusuf AM, Crystal E, et al. Interventions for preventing post-operative atrial fibrillation in patients undergoing heart surgery. Cochrane Database Syst Rev. 2013:CD003611.

13. Larsson SC, Drca N, Michaëlsson K. Serum magnesium and calcium levels and risk of atrial fibrillation. Circ Genom Precis Med. 2019;12:e002349.

14. Nguyen KT, Vittinghoff E, Dewland TA, et al. Ectopy on a single 12-lead ECG, incident cardiac myopathy, and death in the community. J Am Heart Assoc. 2017;6:e006028.

15. Lutsey PL, Chen LY, Eaton A, et al. A pilot randomized trial of oral magnesium supplementation on supraventricular arrhythmias. Nutrients. 2018;10:E884.

16. El-Sherif N, Turitto G. Electrolyte disorders and arrhythmogenesis. Cardiol J. 2011;18:233–245.

17. Landstrom AP, Dobrev D, Wehrens XHT. Calcium signaling and cardiac arrhythmias. Circ Res. 2017;120:1969–1993.

