## Supplementary Results for "Circulating electrolytes and the prevalence of atrial fibrillation and supraventricular ectopy: the Atherosclerosis Risk in Communities (ARIC) Study"

**SUPPLEMENTAL MATERIAL**

**Supplemental Table S1**. Association of circulating electrolytes with prevalence of atrial fibrillation in participants not using diuretics, angiotensin converting enzyme inhibitors, or angiotensin receptor blockers, ARIC study, 2011-2013

| Magnesium | Q1 | Q2 | Q3 | Q4 | Q5 | Per 1-SD^3^ |
| --- | --- | --- | --- | --- | --- | --- |
| Range, mg/dL | 1.2 - 1.9 | 2.0 - 2.0 | 2.1 - 2.1 | 2.2 - 2.2 | 2.3 - 2.7 |  |
| AF | 55 | 37 | 41 | 30 | 14 |  |
| N | 581 | 583 | 675 | 435 | 242 |  |
| OR (95%CI)^1^ | 1 (ref) | 0.62 (0.40, 0.96) | 0.62 (0.40, 0.95) | 0.63 (0.39, 1.01) | 0.52 (0.28, 0.96) | 0.70 (0.59, 0.83) |
| OR (95%CI)^2^ | 1 (ref) | 1.03 (0.74, 1.43) | 1.07 (0.78, 1.47) | 1.02 (0.71, 1.46) | 0.73 (0.45, 1.19) | 0.79 (0.66, 0.94) |
| Calcium | Q1 | Q2 | Q3 | Q4 | Q5 | Per 1-SD^3^ |
| Range, mg/dL | 7.0 - 9.1 | 9.2 - 9.3 | 9.4 - 9.4 | 9.5 - 9.7 | 9.8 - 10.9 |  |
| AF | 72 | 46 | 12 | 31 | 16 |  |
| N | 826 | 612 | 282 | 520 | 276 |  |
| OR (95%CI)^1^ | 1 (ref) | 0.87 (0.59, 1.29) | 0.51 (0.27, 0.96) | 0.68 (0.44, 1.07) | 0.74 (0.41, 1.32) | 0.84 (0.71, 1.00) |
| OR (95%CI)^2^ | 1 (ref) | 1.21 (0.88, 1.66) | 0.71 (0.42, 1.18) | 0.93 (0.64, 1.33) | 0.94 (0.59, 1.51) | 0.84 (0.70, 1.01) |
| Potassium | Q1 | Q2 | Q3 | Q4 | Q5 | Per 1-SD^3^ |
| Range, mmol/L | 2.9 - 3.8 | 3.9 - 4.0 | 4.1 - 4.1 | 4.2 - 4.3 | 4.4 - 6.6 |  |
| AF | 26 | 41 | 31 | 41 | 38 |  |
| N | 487 | 655 | 358 | 583 | 433 |  |
| OR (95%CI)^1^ | 1 (ref) | 0.80 (0.63, 1.03) | 0.90 (0.67, 1.20) | 0.81 (0.63, 1.04) | 0.89 (0.69, 1.15) | 1.28 (1.05, 1.55) |
| OR (95%CI)^2^ | 1 (ref) | 0.99 (0.72, 1.35) | 1.35 (0.95, 1.92) | 0.99 (0.72, 1.36) | 0.95 (0.68, 1.34) | 1.13 (0.92, 1.40) |
| Phosphorus | Q1 | Q2 | Q3 | Q4 | Q5 | Per 1-SD^3^ |
| Range, mg/dL | 1.9 - 3.1 | 3.2 - 3.3 | 3.4 - 3.6 | 3.7 - 3.8 | 3.9 - 6.0 |  |
| AF | 53 | 33 | 45 | 21 | 25 |  |
| N | 606 | 421 | 642 | 366 | 481 |  |
| OR (95%CI)^1^ | 1 (ref) | 0.95 (0.60, 1.53) | 0.92 (0.60, 1.42) | 0.75 (0.43, 1.31) | 0.72 (0.42, 122) | 0.91 (0.75, 1.10) |
| OR (95%CI)^2^ | 1 (ref) | 1.27 (0.90, 1.79) | 1.02 (0.75, 1.39) | 0.88 (0.58, 1.33) | 0.72 (0.49, 1.06) | 0.83 (0.69, 1.01) |
| Chloride | Q1 | Q2 | Q3 | Q4 | Q5 | Per 1-SD^3^ |
| Range, mmol/L | 84 - 98 | 99 - 100 | 101 - 101 | 102 - 103 | 104 - 111 |  |
| AF | 42 | 45 | 14 | 46 | 30 |  |
| N | 389 | 645 | 386 | 682 | 414 |  |
| OR (95%CI)^1^ | 1 (ref) | 0.63 (0.40, 0.99) | 0.33 (0.18, 0.62) | 0.64 (0.41, 1.00) | 0.74 (0.44, 1.23) | 0.86 (0.73, 1.03) |
| OR (95%CI)^2^ | 1 (ref) | 1.01 (0.74, 1.38) | 0.56 (0.35, 0.89) | 1.02 (0.75, 1.39) | 1.14 (0.80, 1.63) | 0.89 (0.74, 1.06) |
| Sodium | Q1 | Q2 | Q3 | Q4 | Q5 | Per 1-SD^3^ |
| Range, mmol/L | 124 - 137 | 138 - 139 | 140 - 140 | 141 - 141 | 142 - 147 |  |
| AF | 50 | 51 | 30 | 17 | 29 |  |
| N | 488 | 862 | 497 | 305 | 364 |  |
| OR (95%CI)^1^ | 1 (ref) | 0.63 (0.41, 0.95) | 0.66 (0.41, 1.07) | 0.61 (0.34, 1.09) | 0.88 (0.54, 1.44) | 0.82 (0.70, 0.96) |
| OR (95%CI)^2^ | 1 (ref) | 0.84 (0.63, 1.13) | 0.99 (0.70, 1.41) | 0.84 (0.53, 1.31) | 1.09 (0.76, 1.58) | 0.82 (0.69, 0.97) |

AF: atrial fibrillation. CI: confidence interval. OR: odds ratio.

^1^Unconditional logistic regression adjusted for age, sex, and race

^2^Unconditional logistic regression adjusted for age, sex, race, center, smoking (never, former, current), physical activity, alcohol intake, systolic blood pressure, diastolic blood pressure, diuretic use, other antihypertensive use, diabetes, heart failure, coronary heart disease, eGFR

^3^One standard deviation is 0.2 mg/dL for magnesium, 0.40 mg/dL for calcium, 0.36 mmol/L for potassium, 0.46 mg/dL for phosphorus, 3.1 mmol/L for chloride, and 2.6 mmol/L for sodium

**Supplemental Table S2**. Association of circulating electrolytes with prevalence of premature atrial contractions in standard 12-lead electrocardiogram in participants not using diuretics, angiotensin converting enzyme inhibitors, or angiotensin receptor blockers, ARIC study, 2011-2013

| Magnesium | Q1 | Q2 | Q3 | Q4 | Q5 | Per 1-SD^3^ |
| --- | --- | --- | --- | --- | --- | --- |
| Range, mg/dL | 1.2 - 1.9 | 2.0 - 2.0 | 2.1 - 2.1 | 2.2 - 2.2 | 2.3 - 2.7 |  |
| PAC present | 65 | 57 | 68 | 60 | 23 |  |
| N | 522 | 545 | 629 | 404 | 225 |  |
| OR (95%CI)^1^ | 1 (ref) | 0.82 (0.56, 1.21) | 0.92 (0.63, 1.33) | 1.24 (0.84, 1.83) | 0.78 (0.47, 1.30) | 0.92 (0.79, 1.07) |
| OR (95%CI)^2^ | 1 (ref) | 0.86 (0.66, 1.12) | 1.00 (0.78, 1.28) | 1.33 (1.01, 1.73) | 0.89 (0.61, 1.29) | 0.97 (0.83, 1.13) |
| Calcium | Q1 | Q2 | Q3 | Q4 | Q5 | Per 1-SD^3^ |
| Range, mg/dL | 7.0 - 9.1 | 9.2 - 9.3 | 9.4 - 9.4 | 9.5 - 9.7 | 9.8 - 10.9 |  |
| PAC present | 92 | 69 | 35 | 48 | 29 |  |
| N | 750 | 564 | 268 | 485 | 258 |  |
| OR (95%CI)^1^ | 1 (ref) | 1.03 (0.73, 1.44) | 1.04 (0.68, 1.60) | 0.73 (0.50, 1.07) | 0.82 (0.51, 1.30) | 0.95 (0.82, 1.09) |
| OR (95%CI)^2^ | 1 (ref) | 1.12 (0.87, 1.43) | 1.16 (0.84, 1.61) | 0.78 (0.59, 1.04) | 0.86 (0.61, 1.23) | 0.92 (0.80, 1.07) |
| Potassium | Q1 | Q2 | Q3 | Q4 | Q5 | Per 1-SD^3^ |
| Range, mmol/L | 2.9 - 3.8 | 3.9 - 4.0 | 4.1 - 4.1 | 4.2 - 4.3 | 4.4 - 6.0 |  |
| PAC present | 46 | 61 | 41 | 69 | 56 |  |
| N | 460 | 611 | 324 | 542 | 388 |  |
| OR (95%CI)^1^ | 1 (ref) | 1.00 (0.66, 1.51) | 1.40 (0.89, 2.22) | 1.37 (0.91, 2.07) | 1.54 (1.00, 2.37) | 1.27 (1.07, 1.50) |
| OR (95%CI)^2^ | 1 (ref) | 0.83 (0.64, 1.07) | 1.15 (0.85, 1.54) | 1.07 (0.84, 1.37) | 1.13 (0.86, 1.49) | 1.18 (0.99, 1.41) |
| Phosphorus | Q1 | Q2 | Q3 | Q4 | Q5 | Per 1-SD^3^ |
| Range, mg/dL | 1.9 - 3.1 | 3.2 - 3.3 | 3.4 - 3.6 | 3.7 - 3.8 | 3.9 - 5.5 |  |
| PAC present | 65 | 42 | 73 | 31 | 62 |  |
| N | 549 | 388 | 594 | 342 | 452 |  |
| OR (95%CI)^1^ | 1 (ref) | 0.84 (0.55, 1.30) | 1.05 (0.72, 1.53) | 0.72 (0.44, 1.15) | 1.22 (0.81, 1.84) | 0.99 (0.85, 1.16) |
| OR (95%CI)^2^ | 1 (ref) | 0.85 (0.63, 1.14) | 1.13 (0.89, 1.44) | 0.75 (0.54, 1.04) | 1.25 (0.96, 1.63) | 0.96 (0.82, 1.13) |
| Chloride | Q1 | Q2 | Q3 | Q4 | Q5 | Per 1-SD^3^ |
| Range, mmol/L | 84 - 98 | 99 - 100 | 101 - 101 | 102 - 103 | 104 - 110 |  |
| PAC present | 46 | 70 | 37 | 86 | 34 |  |
| N | 346 | 599 | 369 | 631 | 380 |  |
| OR (95%CI)^1^ | 1 (ref) | 0.90 (0.60, 1.35) | 0.74 (0.46, 1.18) | 1.09 (0.73, 1.61) | 0.67 (0.41, 1.09) | 0.97 (0.83, 1.12) |
| OR (95%CI)^2^ | 1 (ref) | 1.03 (0.80, 1.31) | 0.87 (0.64, 1.18) | 1.28 (1.01, 1.61) | 0.78 (0.57, 1.07) | 0.98 (0.84, 1.15) |
| Sodium | Q1 | Q2 | Q3 | Q4 | Q5 | Per 1-SD^3^ |
| Range, mmol/L | 124 - 137 | 138 - 139 | 140 - 140 | 141 - 141 | 142 - 147 |  |
| PAC present | 55 | 90 | 60 | 32 | 36 |  |
| N | 436 | 806 | 467 | 287 | 329 |  |
| OR (95%CI)^1^ | 1 (ref) | 0.92 (0.64, 1.33) | 1.06 (0.71, 1.58) | 0.88 (0.55, 1.42) | 0.83 (0.52, 1.31) | 0.99 (0.85, 1.14) |
| OR (95%CI)^2^ | 1 (ref) | 0.99 (0.79, 1.24) | 1.19 (0.92, 1.54) | 0.92 (0.66, 1.28) | 0.84 (0.61, 1.15) | 0.96 (0.83, 1.11) |

CI: confidence interval. OR: odds ratio. PAC: premature atrial contraction

^1^Unconditional logistic regression adjusted for age, sex, and race

^2^Unconditional logistic regression adjusted for age, sex, race, center, smoking (never, former, current), physical activity, alcohol intake, systolic blood pressure, diastolic blood pressure, diuretic use, other antihypertensive use, diabetes, heart failure, coronary heart disease, eGFR

^3^One standard deviation is 0.2 mg/dL for magnesium, 0.40 mg/dL for calcium, 0.36 mmol/L for potassium, 0.46 mg/dL for phosphorus, 3.1 mmol/L for chloride, and 2.6 mmol/L for sodium

**Supplemental Table S3**. Characteristics of participants according to inclusion in the Zio® patch substudy, ARIC study, 2011-2013

|  | | Zio® patch substudy* | | No Zio® patch | |
| --- | --- | --- | --- | --- | --- |
| N | | 317 | | 6073 | |
| Age, years | | 76.9 (5.2) | | 75.7 (5.2) | |
| Women, % | | 53 | | 59 | |
| White, % | | 100 | | 76 | |
| Current smoker, % | | 3.4 | | 6.4 | |
| Alcohol intake, g/week | | 32.6 (65.9) | | 30.9 (64.9) | |
| Body mass index, kg/m^2^ | | 28.3 (4.9) | | 28.8 (5.8) | |
| Hypertension, % | | 80 | | 81 | |
| Diabetes, % | | 32 | | 34 | |
| Heart failure, % | | 7 | | 11 | |
| Coronary heart disease, % | | 13 | | 16 | |
| eGFR, mL/min/1.73 m^2^ | | 66 (16) | | 65 (18) | |
|  | Atrial fibrillation, % | | 9 | | 10 |
| Magnesium, mg/dL | | 2.0 (0.2) | | 2.0 (0.2) | |
|  | Calcium, mg/dL | | 9.3 (0.4) | | 9.4 (0.4) |
| Potassium, mmol/L | | 4.0 (0.3) | | 4.1 (0.4) | |
| Phosphorus, mg/dL | | 3.5 (0.4) | | 3.5 (0.5) | |
|  | Chloride, mmol/L | | 100 (3) | | 100 (3) |
| Sodium, mmol/L | | 138 (3) | | 139 (3) | |

Values correspond to mean (SD) or %. eGFR: estimated glomerular filtration rate.

* Four participants in the Zio® patch study excluded due to recordings <48 hours

**Supplemental Table S4**. Association of circulating electrolytes with burden of premature atrial contractions (PAC), modeled as log_e_(PAC count/day + 1), measured with Zio® patch, ARIC, 2013

| Magnesium | T1 | T2 | T3 | Per 1-SD^4^ |
| --- | --- | --- | --- | --- |
| Range, mg/dL | 1.4 - 1.9 | 2.0 - 2.1 | 2.2 - 2.3 |  |
| PACs / day, mean^1^ | 252 | 330 | 252 |  |
| N | 80 | 132 | 66 |  |
| Beta (95%CI)^2^ | Ref | 0.20 (-0.25, 0.64) | 0.01 (-0.52, 0.53) | 0.06 (-0.14, 0.25) |
| Beta (95%CI)^3^ | Ref | 0.18 (-0.30, 0.66) | -0.05 (-0.62, 0.51) | 0.03 (-0.18, 0.25) |
| Calcium | T1 | T2 | T3 | Per 1-SD^4^ |
| Range, mg/dL | 8.5 - 9.2 | 9.3 - 9.5 | 9.6 - 10.7 |  |
| PACs / day, mean^1^ | 311 | 211 | 395 |  |
| N | 129 | 93 | 56 |  |
| Beta (95%CI)^2^ | Ref | -0.33 (-0.76, 0.10) | 0.15 (-0.37, 0.67) | 0.06 (-0.14, 0.27) |
| Beta (95%CI)^3^ | Ref | -0.27 (-0.71, 0.16) | 0.09 (-0.44, 0.62) | 0.07 (-0.14, 0.28) |
| Potassium | T1 | T2 | T3 | Per 1-SD^4^ |
| Range, mmol/L | 3.0 - 3.9 | 4.0 - 4.2 | 4.3 - 4.9 |  |
| PACs / day, mean^1^ | 308 | 257 | 302 |  |
| N | 105 | 112 | 61 |  |
| Beta (95%CI)^2^ | Ref | -0.25 (-0.67, 0.18) | -0.02 (-0.53, 0.50) | -0.01 (-0.26, 0.23) |
| Beta (95%CI)^3^ | Ref | -0.27 (-0.71, 0.17) | 0.05 (-0.49, 0.60) | 0.00 (-0.26, 0.27) |
| Phosphorus | T1 | T2 | T3 | Per 1-SD^4^ |
| Range, mg/dL | 2.3 - 3.3 | 3.4 - 3.7 | 3.8 - 4.4 |  |
| PACs / day, mean^1^ | 308 | 257 | 302 |  |
| N | 104 | 101 | 73 |  |
| Beta (95%CI)^2^ | Ref | -0.32 (-0.78, 0.14) | -0.09 (-0.62, 0.44) | 0.01 (-0.24, 0.26) |
| Beta (95%CI)^3^ | Ref | -0.38 (-0.85, 0.09) | -0.13 (-0.68, 0.41) | -0.01 (-0.27, 0.25) |
| Chloride | T1 | T2 | T3 | Per 1-SD^4^ |
| Range, mmol/L | 89 - 99 | 100 - 102 | 103 – 107 |  |
| PACs / day, mean^1^ | 299 | 235 | 420 |  |
| N | 100 | 124 | 54 |  |
| Beta (95%CI)^2^ | Ref | -0.22 (-0.65, 0.20) | 0.30 (-0.23, 0.83) | 0.08 (-0.10, 0.27) |
| Beta (95%CI)^3^ | Ref | -0.25 (-0.69, 0.18) | 0.45 (-0.12, 1.02) | 0.12 (-0.08, 0.33) |
| Sodium | T1 | T2 | T3 | Per 1-SD^4^ |
| Range, mmol/L | 128 - 138 | 139 - 140 | 141 – 145 |  |
| PACs / day, mean^1^ | 276 | 247 | 399 |  |
| N | 118 | 102 | 58 |  |
| Beta (95%CI)^2^ | Ref | -0.12 (-0.54, 0.30) | 0.36 (-0.14, 0.86) | 0.09 (-0.10, 0.28) |
| Beta (95%CI)^3^ | Ref | -0.11 (-0.55, 0.33) | 0.37 (-0.14, 0.89) | 0.12 (-0.08, 0.32) |

CI: Confidence interval. PAC: premature atrial contraction.

^1^Geometric mean. ^2^Linear regression adjusted for age and sex. ^3^Linear regression adjusted for age, sex, center, smoking, physical activity, alcohol intake, systolic blood pressure, diastolic blood pressure, diuretic use, ACEI/ARB use, other antihypertensive use, diabetes, heart failure, coronary heart disease, eGFR. ^4^One standard deviation is 0.2 mg/dL for magnesium, 0.40 mg/dL for calcium, 0.36 mmol/L for potassium, 0.46 mg/dL for phosphorus, 3.1 mmol/L for chloride, and 2.6 mmol/L for sodium

**Supplemental Table S5**. Association of circulating electrolytes with prevalence of supraventricular tachycardia, measured with Zio® patch, ARIC, 2013

| Magnesium | T1 | T2 | T3 | Per 1-SD^3^ |
| --- | --- | --- | --- | --- |
| Range, mg/dL | 1.4 - 1.9 | 2.0 - 2.1 | 2.2 - 2.3 |  |
| SVT present | 72 | 123 | 58 |  |
| N | 80 | 132 | 66 |  |
| OR (95%CI)^1^ | 1 (ref) | 1.48 (0.53, 4.12) | 0.82 (0.28, 2.41) | 0.98 (0.63, 1.55) |
| OR (95%CI)^2^ | 1 (ref) | 1.35 (0.72, 2.54) | 0.69 (0.35, 1.36) | 0.84 (0.51, 1.41) |
| Calcium | T1 | T2 | T3 | Per 1-SD^3^ |
| Range, mg/dL | 8.5 - 9.2 | 9.3 - 9.5 | 9.6 - 10.7 |  |
| SVT present | 116 | 84 | 53 |  |
| N | 129 | 93 | 56 |  |
| OR (95%CI)^1^ | 1 (ref) | 0.83 (0.33, 2.10) | 1.16 (0.30, 4.49) | 0.99 (0.61, 1.62) |
| OR (95%CI)^2^ | 1 (ref) | 0.78 (0.38, 1.60) | 1.35 (0.52, 3.48) | 1.14 (0.66, 1.96) |
| Potassium | T1 | T2 | T3 | Per 1-SD^3^ |
| Range, mmol/L | 3.0 - 3.9 | 4.0 - 4.2 | 4.3 - 4.9 |  |
| SVT present | 96 | 103 | 54 |  |
| N | 105 | 112 | 61 |  |
| OR (95%CI)^1^ | 1 (ref) | 1.18 (0.44, 3.18) | 0.87 (0.29, 2.57) | 1.35 (0.78, 2.32) |
| OR (95%CI)^2^ | 1 (ref) | 1.19 (0.64, 2.22) | 0.71 (0.34, 1.48) | 1.30 (0.69, 2.46) |
| Phosphorus | T1 | T2 | T3 | Per 1-SD^3^ |
| Range, mg/dL | 2.3 - 3.3 | 3.4 - 3.7 | 3.8 - 4.4 |  |
| SVT present | 92 | 93 | 68 |  |
| N | 104 | 101 | 73 |  |
| OR (95%CI)^1^ | 1 (ref) | 0.86 (0.32, 2.34) | 0.68 (0.20, 2.30) | 0.88 (0.50, 1.53) |
| OR (95%CI)^2^ | 1 (ref) | 0.98 (0.49, 1.96) | 0.87 (0.38, 1.99) | 0.89 (0.49, 1.63) |
| Chloride | T1 | T2 | T3 | Per 1-SD^3^ |
| Range, mmol/L | 89 - 99 | 100 - 102 | 103 – 107 |  |
| SVT present | 90 | 115 | 48 |  |
| N | 100 | 124 | 54 |  |
| OR (95%CI)^1^ | 1 (ref) | 1.58 (0.60, 4.16) | 0.99 (0.33, 2.99) | 1.03 (0.68, 1.56) |
| OR (95%CI)^2^ | 1 (ref) | 1.42 (0.75, 2.70) | 0.67 (0.30, 1.47) | 0.92 (0.56, 1.52) |
| Sodium | T1 | T2 | T3 | Per 1-SD^3^ |
| Range, mmol/L | 128 - 138 | 139 - 140 | 141 – 145 |  |
| SVT present | 105 | 95 | 53 |  |
| N | 118 | 102 | 58 |  |
| OR (95%CI)^1^ | 1 (ref) | 1.75 (0.65, 4.68) | 1.25 (0.41, 3.81) | 1.03 (0.68, 1.56) |
| OR (95%CI)^2^ | 1 (ref) | 1.14 (0.48, 2.69) | 1.34 (0.49, 3.67) | 0.97 (0.61, 1.54) |

^1^Unconditional logistic regression adjusted for age and sex

^2^Unconditional logistic regression adjusted for age, sex, center, smoking (never v former, current), physical activity, alcohol intake, systolic blood pressure, diastolic blood pressure, diuretic use, ACEI/ARB use, other antihypertensive use, diabetes, heart failure, coronary heart disease, eGFR

^3^One standard deviation is 0.2 mg/dL for magnesium, 0.40 mg/dL for calcium, 0.36 mmol/L for potassium, 0.46 mg/dL for phosphorus, 3.1 mmol/L for chloride, and 2.6 mmol/L for sodium

**Supplemental Table S6**. Association of circulating electrolytes with burden of supraventricular tachycardia (SVT), modeled as log_e_(SVC count/day + 1), measured with Zio® patch, ARIC, 2013

| Magnesium | T1 | T2 | T3 | Per 1-SD^4^ |
| --- | --- | --- | --- | --- |
| Range, mg/dL | 1.4 - 1.9 | 2.0 - 2.1 | 2.2 - 2.3 |  |
| SVT / day, mean^1^ | 1.84 | 2.59 | 2.20 |  |
| N | 80 | 132 | 66 |  |
| Beta (95%CI)^2^ | Ref | 0.32 (0.10, 0.53) | 0.19 (-0.07, 0.44) | 0.09 (-0.00, 0.19) |
| Beta (95%CI)^3^ | Ref | 0.27 (0.04, 0.49) | 0.10 (-0.16, 0.37) | 0.07 (-0.03, 0.17) |
| Calcium | T1 | T2 | T3 | Per 1-SD^4^ |
| Range, mg/dL | 8.5 - 9.2 | 9.3 - 9.5 | 9.6 - 10.7 |  |
| SVT / day, mean^1^ | 2.16 | 2.18 | 2.56 |  |
| N | 129 | 93 | 56 |  |
| Beta (95%CI)^2^ | Ref | -0.01 (-0.22, 0.20) | 0.08 (-0.17, 0.33) | 0.02 (-0.08, 0.12) |
| Beta (95%CI)^3^ | Ref | 0.02 (-0.18, 0.23) | 0.11 (-0.14, 0.37) | 0.06 (-0.04, 0.15) |
| Potassium | T1 | T2 | T3 | Per 1-SD^4^ |
| Range, mmol/L | 3.0 - 3.9 | 4.0 - 4.2 | 4.3 - 4.9 |  |
| SVT / day, mean^1^ | 2.39 | 2.10 | 2.34 |  |
| N | 105 | 112 | 61 |  |
| Beta (95%CI)^2^ | Ref | -0.11 (-0.32, 0.09) | -0.02 (-0.27, 0.23) | -0.00 (-0.12, 0.12) |
| Beta (95%CI)^3^ | Ref | -0.16 (-0.37, 0.05) | -0.06 (-0.32, 0.20) | -0.04 (-0.17, 0.08) |
| Phosphorus | T1 | T2 | T3 | Per 1-SD^4^ |
| Range, mg/dL | 2.3 - 3.3 | 3.4 - 3.7 | 3.8 - 4.4 |  |
| SVT / day, mean^1^ | 2.05 | 2.36 | 2.41 |  |
| N | 104 | 101 | 73 |  |
| Beta (95%CI)^2^ | Ref | 0.05 (-0.18, 0.28) | 0.04 (-0.22, 0.31) | 0.09 (-0.03, 0.21) |
| Beta (95%CI)^3^ | Ref | 0.05 (-0.18, 0.28) | 0.04 (-0.22, 0.31) | 0.09 (-0.03, 0.21) |
| Chloride | T1 | T2 | T3 | Per 1-SD^4^ |
| Range, mmol/L | 89 - 99– | 100 - 102 | 103 – 107 |  |
| SVT / day, mean^1^ | 2.36 | 2.12 | 2.39 |  |
| N | 100 | 124 | 54 |  |
| Beta (95%CI)^2^ | Ref | -0.10 (-0.30, 0.11) | 0.02 (-0.24, 0.28) | 0.00 (-0.09, 0.09) |
| Beta (95%CI)^3^ | Ref | -0.16 (-0.37, 0.05) | -0.00 (-0.28, 0.27) | -0.01 (-0.11, 0.09) |
| Sodium | T1 | T2 | T3 | Per 1-SD^4^ |
| Range, mmol/L | 128 - 138 | 139 - 140 | 141 - 145 |  |
| SVT / day, mean^1^ | 2.20 | 2.20 | 2.44 |  |
| N | 118 | 102 | 58 |  |
| Beta (95%CI)^2^ | Ref | -0.00 (-0.21, 0.21) | 0.09 (-0.16, 0.34) | 0.02 (-0.07, 0.11) |
| Beta (95%CI)^3^ | Ref | -0.04 (-0.25, 0.17) | 0.06 (-0.18, 0.31) | 0.03 (-0.07, 0.12) |

CI: Confidence interval. SVT: supraventricular tachycardia.

^1^Geometric mean. ^2^Linear regression adjusted for age and sex. ^3^Linear regression adjusted for age, sex, center, smoking, physical activity, alcohol intake, systolic blood pressure, diastolic blood pressure, diuretic use, ACEI/ARB use, other antihypertensive use, diabetes, heart failure, coronary heart disease, eGFR. ^4^One standard deviation is 0.2 mg/dL for magnesium, 0.40 mg/dL for calcium, 0.36 mmol/L for potassium, 0.46 mg/dL for phosphorus, 3.1 mmol/L for chloride, and 2.6 mmol/L for sodium
